# Intertissue mechanical interactions shape the olfactory circuit in zebrafish

**DOI:** 10.1101/2021.03.15.435408

**Authors:** P Monnot, G Gangatharan, M Baraban, K Pottin, M Cabrera, I Bonnet, MA Breau

**Author notes:** co-corresponding authors: Marie Anne Breau and Isabelle Bonnet.

## Abstract

While the chemical signals guiding neuronal migration and axon elongation have been extensively studied, the influence of mechanical cues on these processes remains poorly studied *in vivo*. Here, we investigate how mechanical forces exerted by surrounding tissues steer neuronal movements and axon extension during the morphogenesis of the olfactory placode in zebrafish. We mainly focus on the mechanical contribution of the adjacent eye tissue, which develops underneath the placode through extensive evagination and invagination movements. Using quantitative analysis of cell movements and biomechanical manipulations, we show that the developing eye exerts lateral traction forces on the olfactory placode through extracellular matrix, mediating proper morphogenetic movements and axon extension within the placode. Our data shed new light on the key participation of intertissue mechanical interactions in the sculpting of neuronal circuits.

## Introduction

Neuronal circuits develop through a series of dynamic processes including the migration of neurons to their final position and the growth of axons towards target tissues. According to the current view of neuronal development, these processes are primarily guided by chemical cues acting as traffic signs to orient the movement of neurons and of their growing projections^1–3^. In addition to the well-studied chemical signals, developing neurons are also exposed to a variety of mechanical cues, including the stiffness of the environment, and tensile or compressive stresses exerted by surrounding tissues undergoing growth or morphogenesis. Over the past decades, there has been increasing evidence of the influence of such mechanical cues in shaping neuronal morphology *in vitro* (for reviews see references^4,5^). For instance, pulling on neurites can control axon specification, growth, and pruning in neuronal cultures^6–13^. The stiffness of the dish substrate also plays a role in axon growth and branching^14–17^. In contrast, few studies have interrogated the role of such mechanical cues *in vivo*. For example, a brain stiffness gradient influences axon growth patterns in the Xenopus optic pathway^18,19^ and in Drosophila, mechanical tension at the neuromuscular junction modulates vesicle accumulation and synaptic plasticity^20,21^. A better understanding of how mechanical signaling contributes to the construction of neuronal circuits is thus required.

Here, we use the development of a neurogenic placode in zebrafish to investigate how intertissue mechanical interactions influence neuronal movements and axon extension. Neurogenic placodes are transient ectodermal structures contributing sensory neurons to the cranial peripheral nervous system; they include the olfactory, otic, trigeminal and epibranchial placodes^22,23^. In the embryo, the placodes assemble through the coalescence of progenitors located in large domains of the pan-placodal region^24,25^, while or soon before initiating axonal contacts with the brain^26,27^. During their assembly, neurogenic placodes are surrounded by several non-placodal tissues undergoing morphogenetic reorganisation, including the brain^27,28,29^, the optic cup^30^ and neural crest cells^28,31^. The superficial location of the zebrafish neurogenic placodes, right underneath the epidermis, facilitates live imaging of cell dynamics and biomechanical manipulation. This makes them amenable models to explore the influence of pulling/pushing forces from neighbouring tissues on neuronal migration and axon growth and, more globally, the role of intertissue mechanical interactions in morphogenesis, a thriving question in developmental biology^32,33^.

Our study focuses on the morphogenesis of the olfactory placode (OP), which later gives rise to the olfactory epithelium. OP cells are initially located in two elongated domains next to the brain, which both coalesce into paired ellipsoidal neuronal clusters over 7 hours of development, between 14 and 21 hours post fertilization (hpf) (corresponding to 12 somites (12s) and 24s stages)^27,31,34^. We previously showed that OP coalescence is driven by two types of cell movements^27^ (Figure 1A). Cells from the anterior and posterior extremities migrate actively along the brain towards the placode center, and are then displaced laterally, away from the surface of the brain. Central OP cells undergo lateral movements only, without clear anteroposterior displacement. Remarkably, axon growth initiates during lateral movements through retrograde extension, whereby cell bodies move away from their axon tips attached to the brain^27^ (Figure 1A). While the anteroposterior active convergence is guided by Cxcr4b/Cxcl12a signaling downstream of the transcription factor Neurogenin1^29,35^, the mechanisms driving the lateral phase of OP cell movements, during which axons elongate, remain elusive. We previously proposed that the lateral displacement of the cell bodies is a passive, non-autonomous process, based on the following findings: (1) the lack of protrusive activity and of polarised actin and myosin II in the laterally moving cell bodies, (2) the absence of lateral movement defects upon inhibition of microtubule polymerisation or upon cell-autonomous perturbation of actomyosin activity and (3) the lack of lateral movement defects when we disrupted the axons or their attachment to the brain^27^. Altogether, these observations suggest that lateral movements and axon elongation are driven by extrinsic pushing or pulling forces exerted on the OP cell bodies by surrounding cells or tissues.

**Figure 1.**
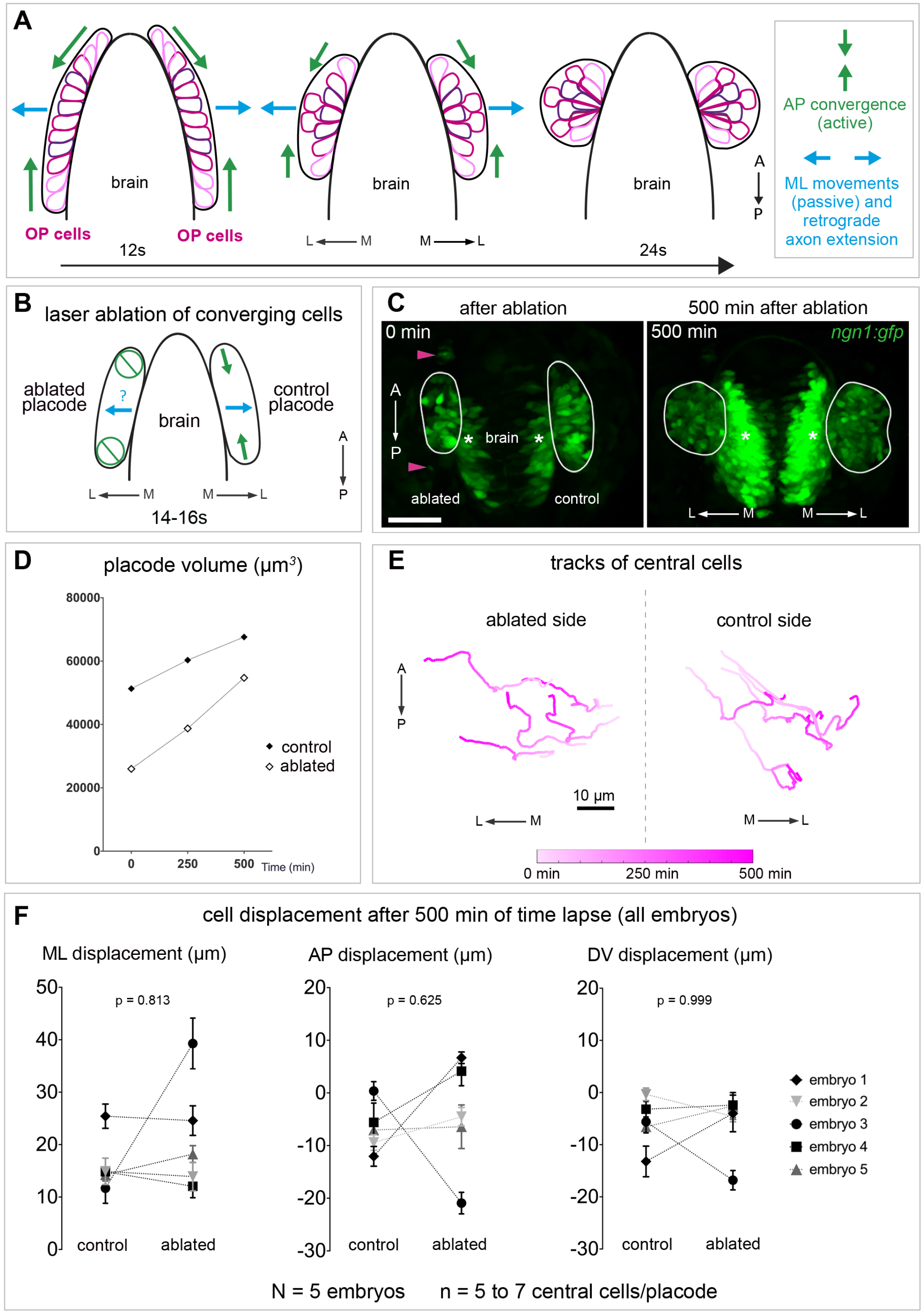
OP cells undergoing anteroposterior convergence are not required for lateral movements. **A**. Schematic view of the two movements driving OP coalescence from 12s (14 hpf) to 24s (21 hpf) (dorsal view, membranes of OP cells in magenta). OP cells are initially located in two elongated domains surrounding the brain. Cells from OP extremities converge towards the centre through active migration along the brain surface (anteroposterior convergence, green arrows), while cell bodies of central cells passively move away from the brain (lateral movements, blue arrows). Axon elongation is concomitant with passive lateral movements and occurs through a retrograde mode of extension, where cell bodies move away from axon tips attached to the brain’s surface. **B**. Schematic view of the laser ablation of converging cells: we used the *ngn1:gfp* transgenic line to ablate the cells undergoing anteroposterior convergence in one of the two placodes (left) at 14-16s, and tracked the central cells during the following 500 min (to reach the 24s stage). Central cells in the contralateral placode were also tracked and used as controls. Dorsal view. **C**. Images of a representative embryo (embryo 1) following the ablation (left) and 500 min later (right) (dorsal view). The ablated (left) and control (right) OPs are surrounded by white lines. Magenta arrowheads show cellular debris where cells were killed in the anterior and posterior regions of the *ngn1:gfp*^+^ placodal cluster. Asterisks indicate gfp expression in the brain. Scale bar: 40 µm. See also Video 1. **D**. Graph showing the volumes of the ablated and control *ngn1:gfp*^+^ placodes for 3 time-points in embryo 1, t = 0 refers to the ablation. **E**. 2D tracks of 5 central cells in the ablated and control placodes in embryo 1. The time is color-coded: light magenta at the starting of the trajectory (just after the ablation) towards dark magenta for the end of the track (500 min later). Similar data (images, volume and tracks) for embryos 2 to 5 are presented in Figure S1. **F**. Graphs showing the total mediolateral (ML), anteroposterior (AP) and dorsoventral (DV) displacements of OP central cells during the 500 min following the ablation, both in ablated and control placodes, for all embryos. N = 5 embryos from 3 independent experiments, each dot represents the mean displacement per placode (+/- sem) calculated from n = 5 to 7 cells/placode, two-tailed Wilcoxon matched-pairs signed rank test.

The goal of this study is to identify the source of the extrinsic mechanical forces involved. We rule out a role for actively converging placodal cells in squeezing out the neuronal cell bodies laterally, and show that the driving force comes from the morphogenesis of the adjacent eye tissue, transmitted by mutual physical interactions with the intervening extracellular matrix (ECM). This work sheds new light on the role of mechanical forces exchanged between developing neurons and surrounding tissues in the sculpting of neuronal circuits *in vivo*, which was largely unexplored so far.

## Results

### OP cells undergoing anteroposterior convergence are not required for OP lateral movements

We initially hypothesised that cells from the anterior and posterior extremities of the OP -while actively converging -may compress the central cells, thereby squeezing them away from the brain and contributing to the elongation of their axons^27^. To test this, we ablated the cells that would exert this compression, using the *Tg(−8*.*4neurog1:GFP)* line^36^ referred to below as the *ngn1:gfp* line, which labels the early-born neurons in the OP^27,37^. In practise, we laser ablated 10 to 20 *ngn1:gfp*^+^ cells in both anterior and posterior extremities of one placode when convergence occurs, at 14-16s (the placode typically comprising a hundred *ngn1:gfp*^+^ cells at this stage^27^), the contralateral placode serving as a control (Figure 1B,C). We then performed live imaging and 3D cell tracking of the central cells in the ablated placode, and compared their movements with those of central cells in the control placode. We focused on embryos in which ablated placodes displayed smaller volumes than contralateral placodes throughout OP morphogenesis, over 500 min after the ablation, indicating a successful ablation (N = 5 embryos; Figure 1C,D, Figure S1A,B and Video 1). No significant defect in the mediolateral displacement of central OP cells was detected in the ablated placode (nor in the anteroposterior or dorsoventral displacements, Figure 1E,F, Figure S1C and Video 1). These results rule out a major contribution of actively converging OP cells in mediating the lateral movements of central cells. We thus turned our attention to another potential source of extrinsic mechanical forces: the adjacent eye tissue.

### OP and eye cell movements are correlated along the mediolateral axis

The optic cup develops close to the OP in a deeper, more ventral position, through extensive evagination and invagination movements having both a strong lateral component ^30,38–44^ (Figure 2A). Such morphogenetic processes may exert forces on the overlying OP neurons and contribute to their lateral movement and axon extension. To test the influence of optic cup morphogenesis on OP formation, we first compared cell movements in the OP with those occurring in the forming eye, focusing on their lateral component. To do so, we performed live imaging using a frontal view, on embryos carrying the *ngn1:gfp* transgene, labelling the early-born neurons in the OP but also a subpopulation of neurons in the anterior brain^27^, and the *Tg(rx3:GFP)* transgene (*rx3:gfp*), expressed by neural retina progenitors^45^ (N = 4 embryos, Video 2). We used a biological reference time Tref, which represents the time when the eye invagination angle is equal to 120°, to synchronise the 4 embryos and analyse a common time-window around Tref (see methods and Figure S2A,B). Taking advantage of nucleus red labelling obtained with H2B-RFP mRNA injection, we tracked individual *ngn1:gfp*^+^ OP cells throughout OP coalescence, from 12s to 24s, and found that they recapitulate the lateral movements we previously described on dorsal views^27^ (Figure 2B and Figure S2E). To analyse morphogenetic movements in the forming optic cup, we focused on *rx3:gfp*^+^ eye cells populating the anterior neural retina, as their movements were shown to take place in the vicinity of the OP, as opposed to other neural retina progenitors^30,40,42,46^. Strikingly, anterior neural retina progenitors displayed mediolateral directional movements that appeared to be coordinated with those of OP cells (Figure 2B and Figure S2E). To evaluate the contribution of other surrounding tissues, we also tracked *ngn1:gfp*^+^ cells in the adjacent forebrain, and skin (peridermal) cells overlying the OP and the eye. Consistent with previous findings^27,29^, skin cells showed spatially limited, non-oriented displacements. Brain *ngn1:gfp*^+^ cells did not show directed lateral movements but often moved back and forth along the mediolateral axis, akin to interkinetic nuclear migration within the brain neuroepithelium (Figure 2B and Figure S2E).

**Figure 2.**
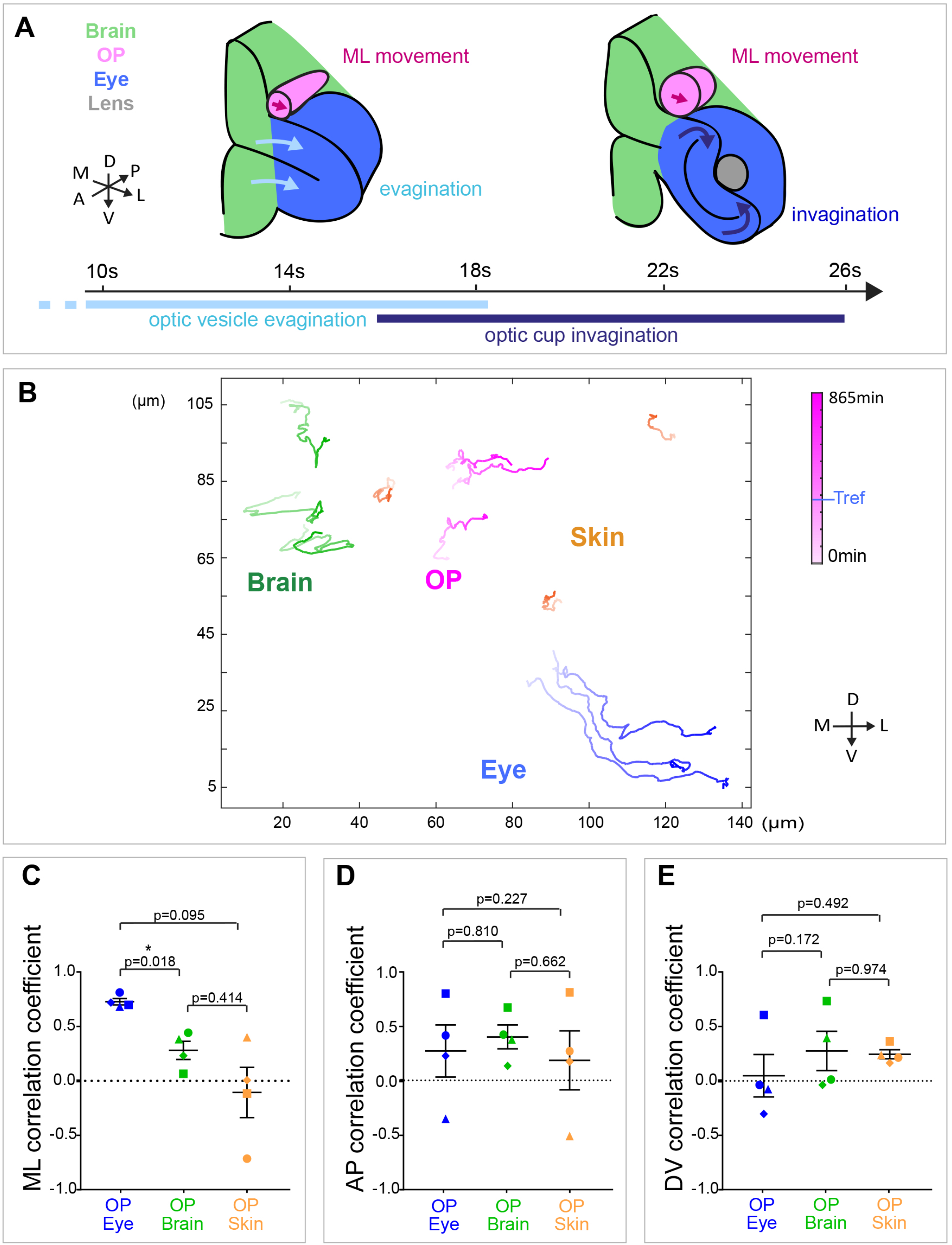
OP and eye cell movements are correlated along the mediolateral axis. **A**. 3D schematic view of optic vesicle evagination and optic cup invagination, occurring during OP coalescence. The optic cup (blue) develops near the OP (pink) in a deeper, more ventral position. The brain is represented in green. Both evagination and invagination movements display a strong lateral component and may exert forces on the overlying OP neurons, contributing to their lateral movement and axon extension. **B**. Live imaging in a frontal view on *ngn1:gfp*; *rx3:gfp* double transgenic embryos injected with H2B:RFP mRNA to label cell nuclei allowed us to track individual cells in four tissues from 12s to 24s (N = 4 embryos from 2 independent experiments, the way we synchronised the 4 movies is detailed in the methods). Three representative trajectories of *ngn1:gfp*^+^ OP cells (magenta), adjacent *ngn1:gfp*^+^ brain cells (green), *rx3:gfp*^*+*^ eye cells populating the anterior retina (blue) and overlying skin cells (orange) in embryo 1. The time is color-coded: light at the starting of the trajectory (12s) towards dark for the end of the track (24s, 865 min later). The reference time (Tref) when the eye invagination angle is 120° (see methods) is shown in the colorbar indicating the time. **C-E**. Mediolateral (ML), anteroposterior (AP) and dorsoventral (DV) correlation between pairs of cells belonging to different tissues. A value of 1 indicates highly correlated tracks and -1 is for anticorrelated tracks. Each dot represents the average correlation coefficient for a given pair of tissues in a given embryo (N = 4 embryos from 2 independent experiments), calculated from the tracking data of 5 to 7 cells per tissue. One-way paired ANOVA with Greenhouse-Geisser correction and Tukey’s multiple comparison test. The correlation coefficients obtained of all pairs of cell trajectories for all embryos are presented in correlation matrices shown in Figure S2F.

To further compare the movements of cells from different tissues, we computed correlation coefficients between pairs of tracks from cells belonging to the four analysed tissues (OP, eye, brain, skin). For each pair of cells, we computed the correlation coefficient of their two trajectories for a given dimension (mediolateral, anteroposterior or dorsoventral component). Correlation coefficients are dimensionless and range from -1 (anticorrelated) to +1 (correlated). Averaging correlation coefficients for pairs of cells from the same tissue reflects the cohesiveness of cell motion in this tissue, while averaging these coefficients for pairs of cells from two different tissues reflects the degree to which the motions of these tissues are correlated along the dimension of interest (see methods). In all embryos, the mean mediolateral correlation coefficient for the eye/OP tissue couple was higher than 0.7 (Figure 2C; Figure S2F), close to values of OP and eye intratissue correlations (Figure S2C), showing that eye cells populating the anterior neural retina and OP cells have coordinated displacements in the mediolateral direction. This high correlation was not solely due to the proximity between the two tissues, since the mediolateral correlation for brain/OP and skin/OP couples was lower than for the eye/OP combination (Figure 2C; Figure S2F). In contrast, the correlation coefficients for the anteroposterior and dorsoventral components of cell trajectories were similar for all tissue pairs, and lower than the eye/OP correlation in the mediolateral axis (Figure 2D,E). The high correlation between eye and OP lateral movements supports the idea of a coupling between the two adjacent tissues. We thus wanted to use perturbative approaches to explore the mechanical interplay between the eye and the OP.

### OP lateral movements are reduced in eyeless *rx3* mutant embryos

To analyse whether eye morphogenesis is required for OP coalescence, we used a mutant for *rx3* (*rx3*^*s399*^), a transcription factor specifically expressed in eye progenitors and the anterior hypothalamus from 8 hpf, and shown to be crucial for eye development^47–53^. While embryos that are heterozygous for this mutation do not show any phenotype, the eye field progenitors in *rx3*^*-/-*^ homozygous embryos do not evaginate to form the optic vesicles, resulting in a complete loss of optic cups^50^. We analysed the volume, the dimensions and the number of cells of the OP, in *rx3*^*-/-*^ mutants versus control siblings at the end of OP morphogenesis (24s). *rx3*^*-/-*^ OPs displayed normal volume and cell numbers (Figure S3). The anteroposterior and dorsoventral dimensions of the placode were increased in *rx3*^*-/-*^ mutants, while the mediolateral dimension was decreased, as compared with control siblings (Figure 3A-C). To assess potential defects in the movements of OP cells during coalescence, we followed OP cells using live imaging in a dorsal view (Video 3) and 3D tracking (N = 5 mutant placodes and N = 3 control placodes). No significant change in the total anteroposterior displacements was detected (Figure 4 and Figure S4A). The dorsoventral movements were not affected either (Figure S4B). By contrast, the lateral displacement of OP cells was significantly reduced in *rx3*^*-/-*^ mutants, while there was no change in the lateral displacement of adjacent *ngn1:gfp*^*+*^ brain cells or overlying skin cells (Figure 4C). Thus, the absence of eyes in *rx3*^*-/-*^ mutants mostly affects the lateral movements of OP cells. Altogether, these results show that optic cup formation is required for proper OP morphogenesis along the mediolateral axis.

**Figure 3.**
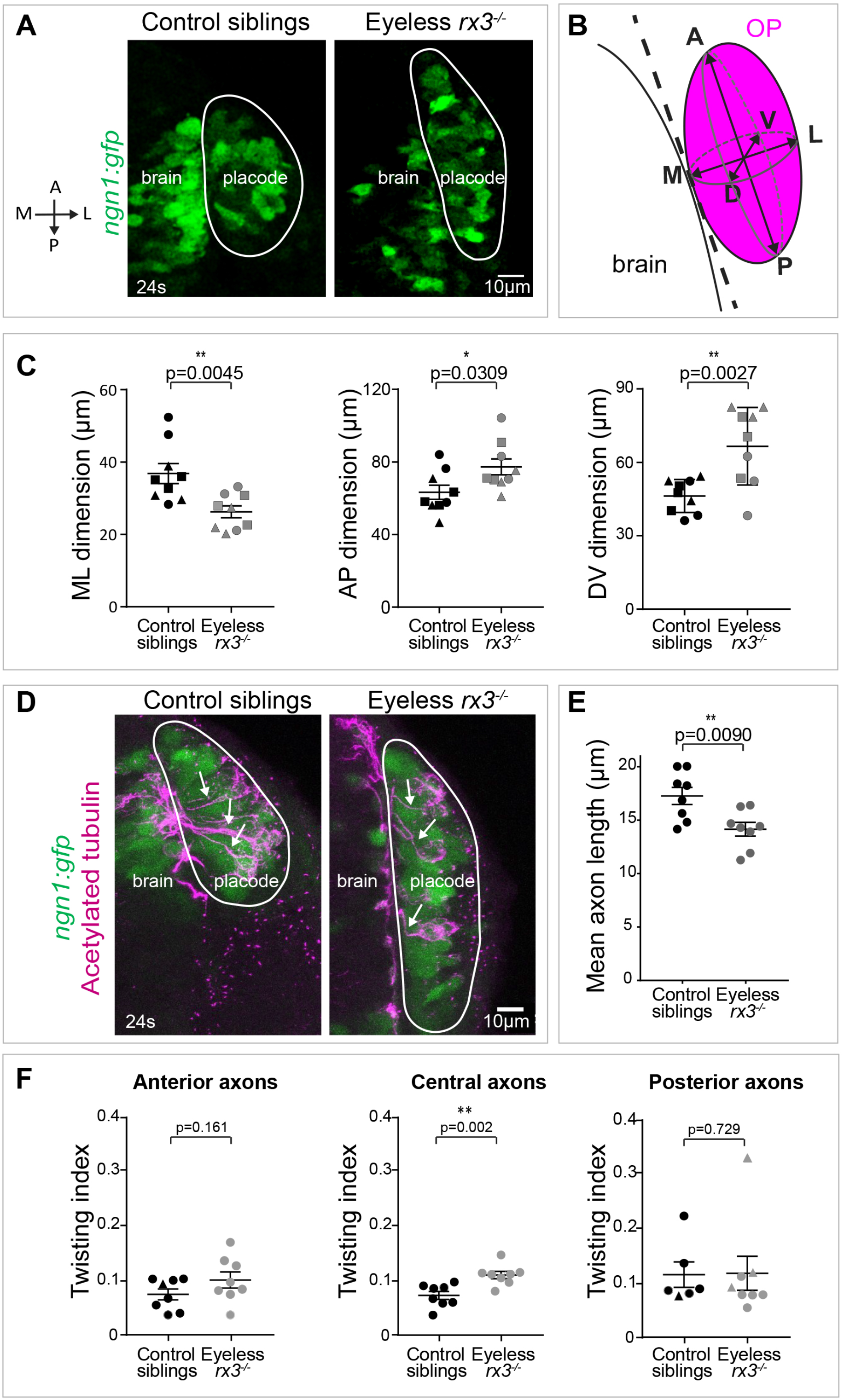
OP shape and axon length are affected in *rx3*^*-/-*^ eyeless embryos. **A**. Images (dorsal views, 1 z-section) of representative placodes from a *ngn1:gfp; rx3*^*-/-*^ mutant (right) and a control *ngn1:gfp* sibling (left) at the end of OP coalescence (24s). The *ngn1:gfp*^+^ OP clusters are surrounded by white lines. **B**. The *ngn1:gfp*^+^ OP clusters were fitted with a 3D ellipsoid (pink) to estimate the mediolateral (ML), anteroposterior (AP), and dorsoventral (DV) dimensions of the placode (details in methods) at 24s. **C**. Graphs showing the mediolateral (ML), anteroposterior (AP), and dorsoventral (DV) dimensions of the OP in *rx3*^*-/-*^ eyeless mutants (grey) and control siblings (black) at 24s (N = 9 embryos from 3 independent experiments). Embryos from the same experiment are represented with similar markers (dots, triangles or squares). Unpaired, two-tailed t test. **D**. Confocal images (dorsal views, 20 µm maximum projections) of representative placodes from a *ngn1:gfp; rx3*^*-/-*^ eyeless mutant (right) and a control sibling (left) immunostained at 24s for acetyl-tubulin (magenta), which labels a subpopulation of *ngn1:gfp*^+^ (green) neurons and their axons. The *ngn1:gfp*^+^ OP clusters are surrounded by white lines. Some axons are indicated by arrows. **E**. Graph showing the length of acetyl-tubulin^+^ axons in *rx3*^*-/-*^ mutants (grey) and control siblings (black) at 24s (N = 8 embryos from 3 independent experiments). Unpaired, two-tailed t test. **F**. Graphs showing the twisting index (see methods) of anterior, central and posterior axons in *rx3*^*-/-*^ mutants (grey) and control siblings (black) at 24s (N = 8 embryos from 3 independent experiments). For embryos represented by a triangle, only one axon was measured. Unpaired, two-tailed t test for anterior and central axons. For the posterior axons the data did not show a normal distribution and a two-tailed Mann-Whitney test was performed.

**Figure 4.**
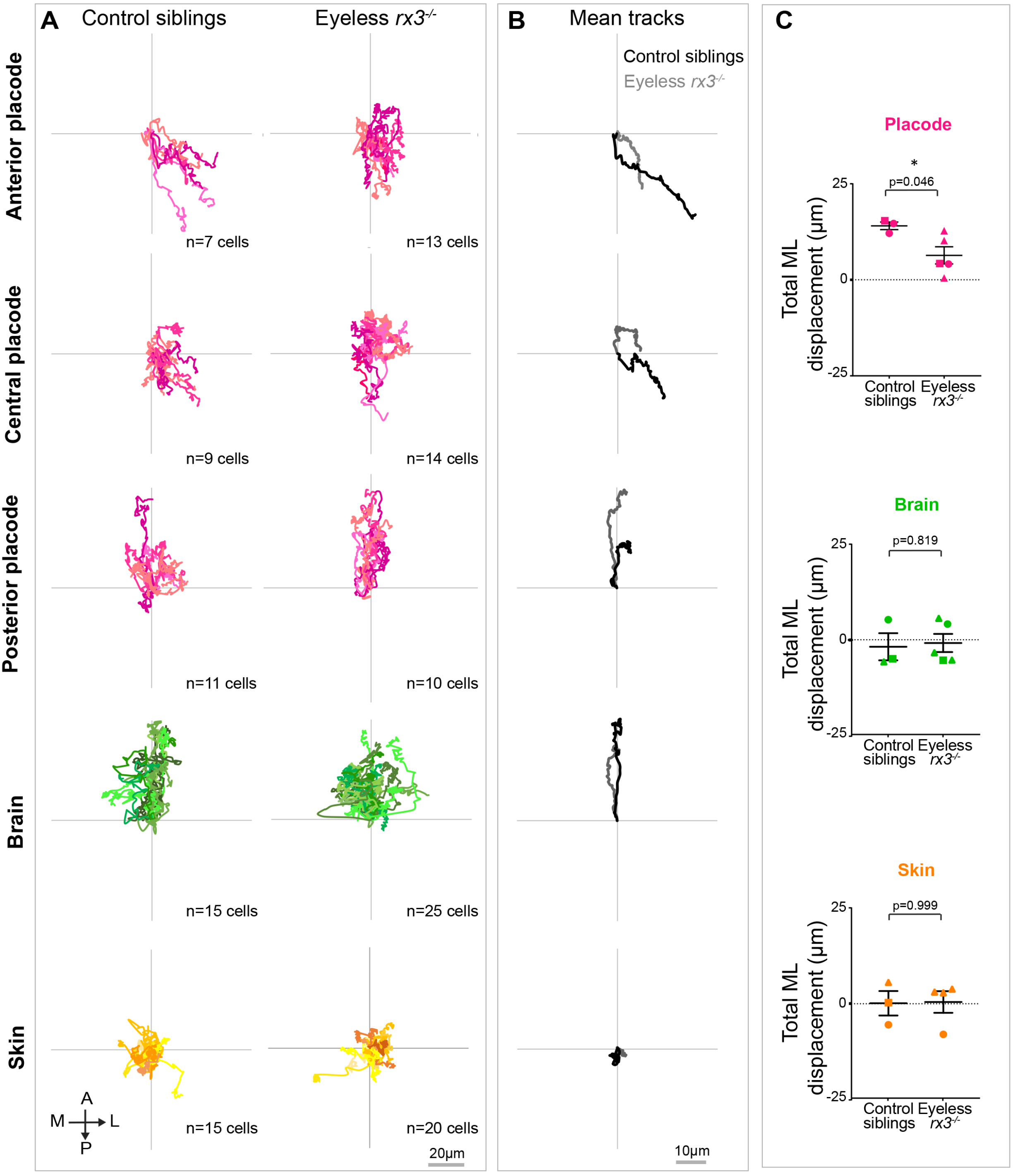
Mediolateral movements of OP cells are reduced in *rx3*^*-/-*^ eyeless embryos. **A**. 2D tracks (mediolateral along X and anteroposterior along Y), merged at their origin, of anterior, central and posterior *ngn1:gfp*^+^ placodal cells (as defined in^27^) (magenta), *ngn1:gfp*^+^ brain cells (green), and skin cells (orange), in control and *rx3*^*-/-*^ eyeless mutant embryos (N = 3 control placodes and N = 5 mutant placodes from 3 independent experiments). All cells were tracked throughout the morphogenesis process, from 12s to 24s stages, during a 700 min period of time. **B**. Mean 2D trajectories for control and *rx3*^*-/-*^ eyeless mutant embryos. **C**. Graphs showing the total mediolateral (ML) displacement of OP (magenta), brain (green) and skin (orange) cells, starting at 12s and during 700 min of time lapse in *rx3*^*-/-*^ eyeless mutants and control embryos (N = 3 control placodes and N = 5 mutant placodes from 3 independent experiments, 5 to 7 cells per tissue per placode, unpaired, two-tailed t test).

### Axons in the OP are shorter and more twisted in eyeless *rx3* mutant embryos

Since we previously showed that axon growth initiates during lateral movement through retrograde extension^27^, we also wanted to assess the influence of optic cup morphogenesis on axon elongation. To this end, we analysed the length and shape of axons in *rx3*^*-/-*^ mutant embryos at the end of OP coalescence (24s, Figure 3D). Consistent with a reduced mediolateral dimension of the OP and decreased lateral movements of OP cells, the axons of acetylated-tubulin^+^ neurons (a subpopulation of *ngn1:gfp*^*+*^ neurons) were shorter in *rx3*^*-/-*^ embryos in comparison to control siblings (Figure 3E). These results show that optic cup morphogenesis also contributes to the retrograde elongation of axons within the OP. Interestingly, we also observed that axons located in the central region (along the anteroposterior axis) of the OP tissue, that elongate the most parallel to the mediolateral orientation, were more twisted in *rx3*^*-/-*^ embryos than in controls (Figure 3F). These observations suggest that, in addition to being shorter, axons are less stretched along the mediolateral axis in the absence of optic cups. At this point, all our findings support the idea of a mechanical coupling between the eye and the OP, whereby the forming eye pulls laterally on neuronal cell bodies and their trailing axons.

### Mechanical tension at the lateral border of the OP is decreased in eyeless *rx3* mutant embryos

To further probe this mechanical coupling, we used linear laser ablations. Laser ablations were initially used to sever subcellular structures that support force transmission to provoke a sudden mechanical imbalance: the initial recoil velocity of the ablated structure gives, under some assumptions, an estimation of the tension prior ablation^54–56^. Laser ablation can also be performed at tissue-scale to create a wound and estimate tissue stress^57–61^. To estimate mechanical stresses at the periphery of the *ngn1:gfp*^+^ cell group within the OP, we performed linear supracellular laser cuts in anterior, lateral and posterior borders of the *ngn1:gfp*^+^ cluster in control embryos and *rx3*^*-/-*^ mutants, at 16s (as depicted in Figure 5A, Video 4). We analysed tissue relaxation using particle image velocimetry (Figure 5B). Assuming that the tissue material properties are homogenous, the initial retraction velocity gives a relative estimation of the tissue mechanical stress before the cut. Mechanical stress was significantly reduced at the lateral border of *rx3*^*-/-*^ mutants compared with the controls, whereas no change was detected in anterior and posterior OP borders (Figure 5C). These results show that the tension to which the *ngn1:gfp*^+^ placodal cluster is subjected along the mediolateral axis is reduced when the eye is absent, in agreement with the hypothesis that the developing eye exerts lateral traction forces on OP neurons during OP morphogenesis.

**Figure 5.**
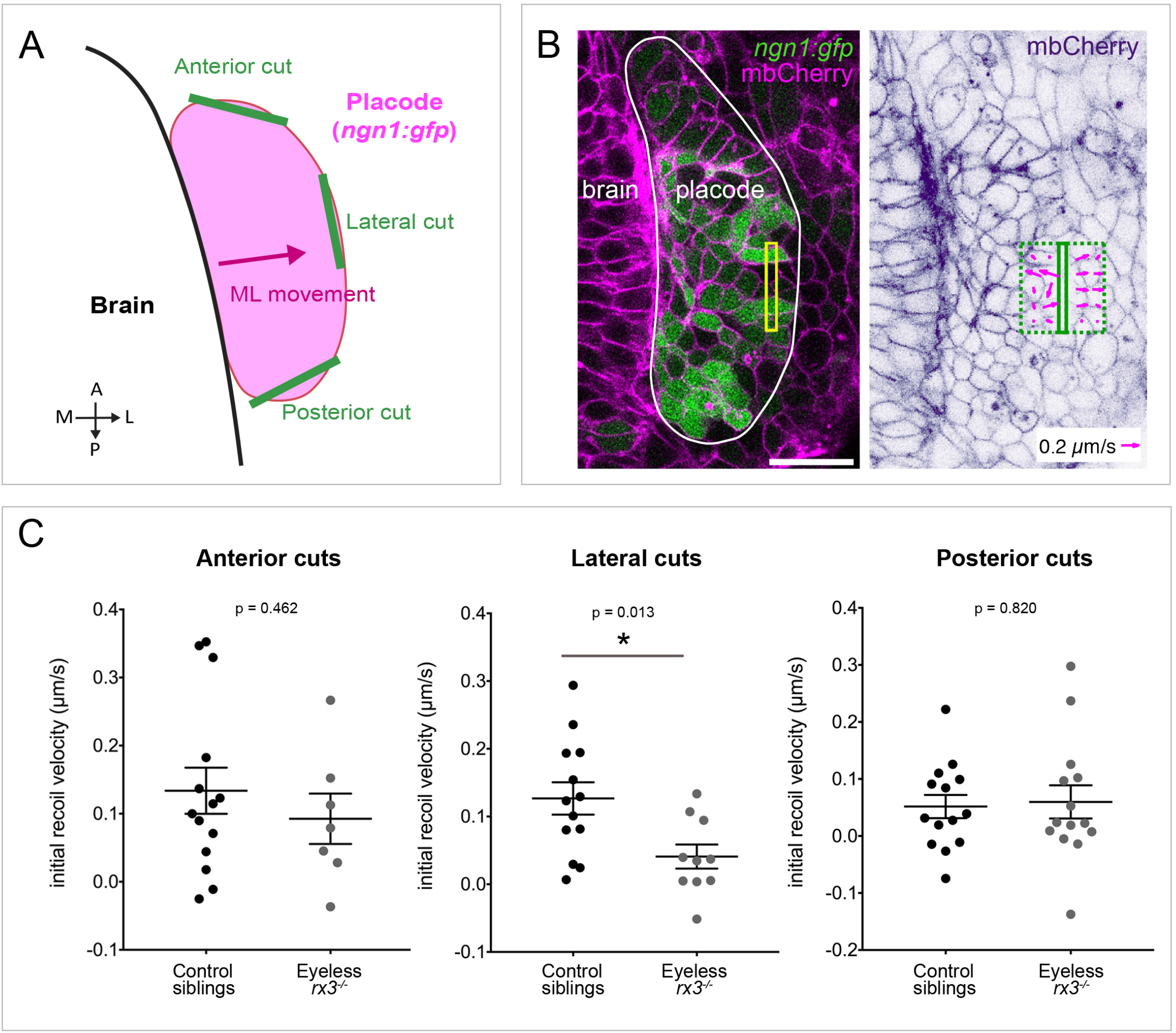
Mechanical stress at the lateral border of the *ngn1:gfp*^+^ OP cluster is reduced in eyeless embryos. **A**. Schematic view of the supracellular laser ablation experiment (dorsal view). Linear laser cuts (green segments) were performed at the anterior, lateral or posterior border of the *ngn1:gfp*^+^ OP cluster at 16s, in control and *rx3*^*-/-*^ embryos injected with mbCherry mRNA to label the membranes. **B**. Example of a lateral cut performed on a control *ngn1:gfp* (green) embryo expressing mbCherry (magenta on the left panel, and purple on the right panel). Left: image showing the position of the region of ablation (yellow), just before the ablation, the OP is delineated with a white line. Scale bar: 20 µm. Right: same image 3 s after the ablation, with tissue velocity field overimposed (magenta arrows) in rectangles of 10 µm width on both sides of the ablation line. **C**. Graphs showing the initial relaxation velocity, used as a proxy for tissue mechanical stress before the cut, following anterior, lateral or posterior cuts, in control and *rx3*^*-/-*^ embryos (data pooled from 7 independent experiments). Unpaired, two-tailed t test.

### Neural crest cells populate the eye/OP interface but are not essential for OP morphogenesis

If the eye pulls on the OP, forces must be transmitted through a physical contact, we thus decided to inspect the interface between these two tissues. To analyse whether the OP and the eye interact through direct intercellular contacts, we took advantage of the *Tg(−8*.*0cldnb:LY-EGFP)* line (*cldnb:lyn-gfp*), in which a membrane-targeted version of gfp is expressed in forebrain, skin (peridermal) cells, but also eye cells (in the neural retina and the retinal pigmented epithelium) and all cells of the OP^62,63^. Imaging the eye/OP interface on *cldnb:lyn-gfp* embryos showed the presence of a gfp negative gap between the two tissues, about 10-15 µm-large, demonstrating the absence of common intercellular junctions (Figure S5A and Videos 5 and 6). Interestingly, we observed the presence of gfp negative cells progressively populating the gap between the two tissues from around 18-20s onwards (Figure S5A). Based on the literature^28,31,46,64^, we hypothesised that these cells are cranial neural crest cells (NCCs). NCCs are known to influence optic cup formation^46^ and their migration is affected in *rx3* mutants^64^. To test whether NCCs are necessary for OP formation, we searched for a condition in which the migration of NCCs populating the eye/OP interface is affected, but not eye morphogenesis. This situation can be found in *foxd3*^-/-^ mutants, as described in a recent study^46^ and confirmed by our analysis of NCCs (Figure S5B) and eye size and invagination in *foxd3*^-/-^ mutants (Figure S5D). The number of cells located in the gap between the two tissues was reduced in *foxd3*^-/-^ mutants, confirming that these interstitial cells mostly represent migrating NCCs (Figure S5E). The volume of the OP, as well as its anteroposterior and dorsoventral dimensions, were not affected in *foxd3*^-/-^ mutants, but the mediolateral dimension was increased (Figure S5D), showing that the NCCs are not required for, but rather prevent OP lateral movements. These results argue against a major role of these interstitial NCCs in mediating OP lateral movements.

### The extracellular matrix establishes a physical link between the eye and the OP

From our experiments, we conclude that the OP and the eye should be tightly linked. To further identify the mechanisms underlying such mechanical coupling, we hypothesized that the two tissues are connected by and interact through ECM. We thus analysed by immunostaining the localisation of two ECM components, Laminin and Fibronectin, at the tissue interface. Laminin displayed a basement membrane-like distribution around the OP and the eye, suggesting the presence of two basement membranes surrounding the eye and the placode respectively (Figure 6A and Figure S5B). Fibronectin showed a more interstitial distribution surrounding the cells located at the interface between the eye and the OP (Figure 6A). These expression patterns were unchanged in *rx3*^*-/-*^ mutants, except the basement membrane surrounding the forming eye, due to the lack of optic cup (Figure 6A), suggesting that the ECM surrounding the OP is not sufficient by itself to drive proper OP morphogenesis. To test whether ECM acts as a glue physically transmitting forces between the eye and the OP, we degraded the ECM by injecting a mix of collagenases and red fluorescent dextran, as previously described in Tlili *et al*. 2020^65^, close to the eye/OP interface of *ngn1:gfp; rx3:gfp* embryos. This resulted in the rapid diffusion of the fluorescent mix in all the extracellular space of the head region (Figure S6A), and in a global perturbation of Laminin distribution (weaker, more diffuse and less continuous signal) at 24s (Figure S6B), confirming the efficiency of the treatment. This approach thus provides a temporal control on ECM perturbation, but no precise spatial control. To specifically target the lateral movements in the OP, and not the anteroposterior convergence movements, we performed the injection at 16-18s, when most of the convergence has already occurred^29^. As expected from previous studies reporting a role for the ECM in optic cup morphogenesis^30,42,66^, the collagenases injection resulted in a range of eye defects (from little or no apparent defect to strong invagination phenotypes often associated with extrusion of the lens), likely reflecting variations in the injected volume, a parameter we can not unfortunately finely tune. In order to address the role of the ECM independently from that of the eye, we selected the collagenases-injected embryos in which eye cells still displayed coherent (with a high intra-eye correlation coefficient) and directional lateral movements (Figure 6B,C and Figure S6C,D). Strikingly, in these embryos, the lateral movements of OP cells were perturbed, and their correlation with the lateral movements of eye cells was decreased, as compared with control embryos injected with dextran only (N = 3 collagenases-injected and N = 3 dextran-injected control embryos) (Figure 6B,C and Figure S6C,D). These data provide evidence that the ECM mechanically couples OP and eye morphogenesis.

**Figure 6.**
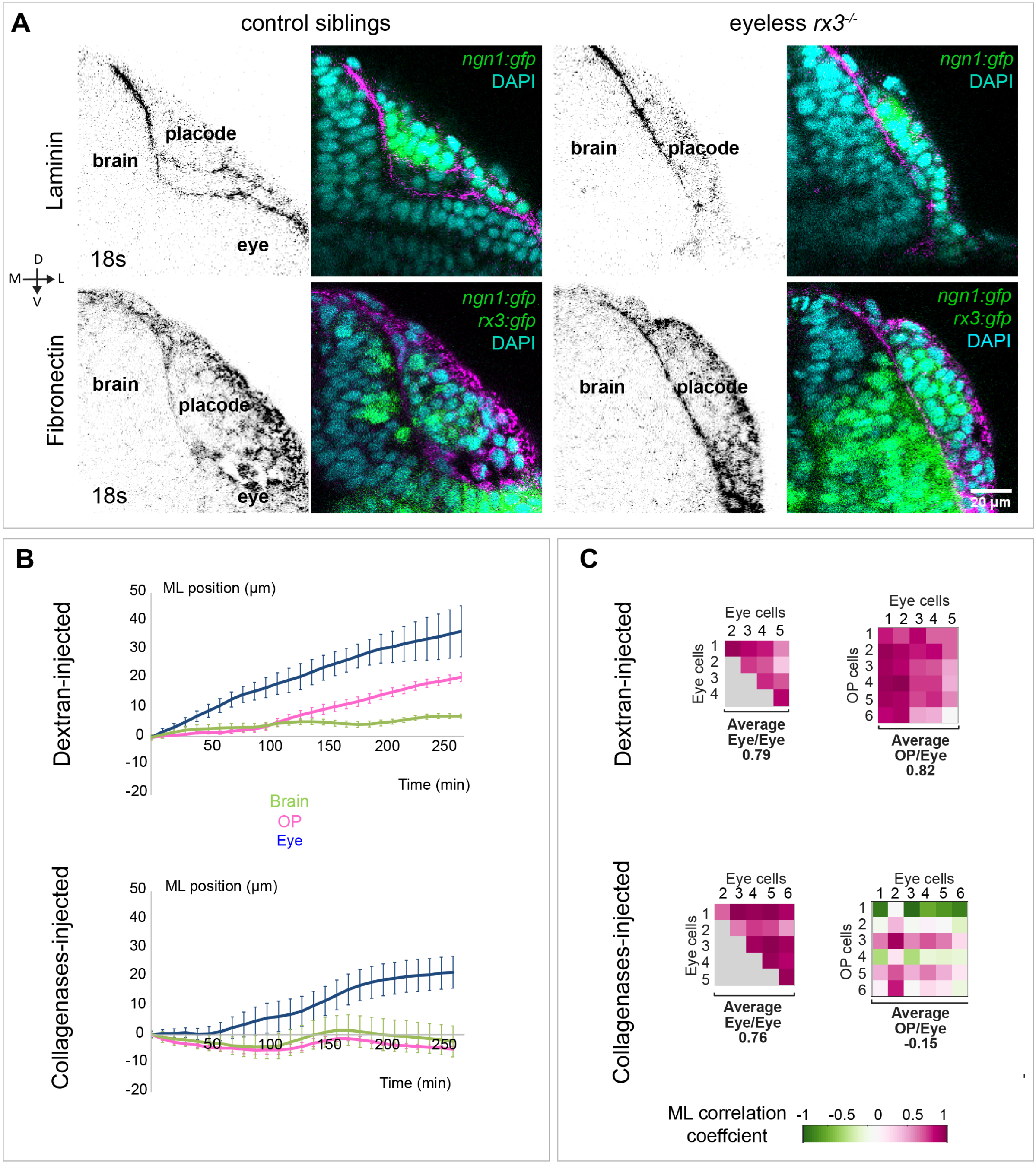
The extracellular matrix mechanically couples the eye to the OP. **A**. Confocal images (frontal views, 1 z-section) of Laminin and Fibronectin immunostainings (black, or magenta on the merge) performed on *ngn1:gfp; rx3:gfp* control (left) and *rx3*^*-/-*^ (right) embryos fixed at 18s. **B**. Live imaging with a dorsal view was performed on *ngn1:gfp*; *rx3:gfp* embryos expressing H2B:RFP and injected at 16s with a mixture of collagenases and red dextran or with dextran only. Individual cells were tracked in several tissues (OP, eye, brain and skin) from 16s to 24s (N = 3 collagenases-injected embryos and N = 3 dextran-injected control embryos from 3 independent experiments). To compensate for the drift often observed in injected embryos, the average displacement of skin cells was used for registration and removed from the tracking data. The graphs show the mediolateral displacement of OP (magenta), eye (blue) and brain (green) cells as a function of time (mean +/- sem from 5 to 7 cells per tissue) for one control and one collagenase-injected embryo (experiment 1). Results for experiments 2 and 3 are shown in Figure S6C. **C**. Correlation matrices presenting the intra-eye and the eye/OP mediolateral correlation coefficients, calculated from the tracking data of 5 to 7 cells per tissue (experiment 1). The average correlation coefficient for each tissue couple is indicated below the matrices. Results for experiments 2 and 3 are shown in Figure S6D.

## Discussion

Our study reveals that the developing optic cup contributes, from a mechanical point of view, to the sculpting of the OP. Our data show that eye morphogenesis steers OP lateral movements and axon extension by exerting pulling forces in the lateral direction, transmitted to OP cells by ECM located at the interface between the two tissues.

The coalescence of the OP is driven by two types of cell movements, anteroposterior convergence and lateral movement^27^. Cxcr4b/Cxcl12a chemotactic signaling has been shown to control OP coalescence downstream of the Neurogenin1 transcription factor^29,35^. While the convergence of the anterior OP cells is particularly affected in mutants for the Cxcr4b/Cxcl12a pathway, the mediolateral component of the cell movements appears to be less perturbed^29^. Conversely, the lateral movements were decreased in eyeless *rx3*^*-/-*^ mutants, but we did not detect significant defects in anteroposterior convergence movements, although the anteroposterior OP dimension was increased at the end of OP coalescence (24s). This increase could be due to a weak perturbation of Cxcl12a/Cxcr4b signaling in *rx3*^*-/-*^ mutants, consistent with the previously described posterior expansion of the telencephalon^67,68^, known to express the Cxcl12a ligand^29,35^. Alternatively, the higher anteroposterior OP dimension in *rx3*^*-/-*^ mutants could be a consequence of the lateral movement defects: because of the inability of OP cells to be displaced laterally by the forming eye, the converging cells would accumulate at the anterior and posterior borders of the OP, thus increasing the dimension of the OP along the anteroposterior axis. Future investigations will distinguish between these hypotheses and clarify how Cxcr4b/Cxcl12a chemotactic signaling and eye mechanical traction cooperate to orchestrate the cell movements that shape the OP.

Eye morphogenesis is a complex, multi-step process, including optic vesicle evagination, optic cup invagination, lens formation and spreading of the retinal pigmented epithelium over the forming cup ^43,44^. Which of these processes is the driving force of OP morphogenesis? In *rx3*^-/-^ mutants, in which OP lateral movements are affected, evagination does not occur but a lens assembles^38^, suggesting that lens formation is not sufficient to drive OP morphogenesis. Evagination of the optic vesicle is proposed to be driven by active cell migration in medaka^38^ and, in zebrafish, by intercalation of cells from the core of the eye field into the retinal neuroepithelium^39^. This results in a massive lateral and posterior cell flow which could act as a conveying belt dragging the overlying OP cells laterally during the early phases of OP coalescence. Later, the invagination i.e. the remodelling of the vesicle into a cup is suggested to occur through basal constriction in the centre of the retina epithelium, which may transmit tension at the tissue scale to trigger the bending of the retina^41,42^. Rim movements of cells from the medial layer of the cup into the lateral layer^40,42^, and the spreading of the retinal pigmented epithelium over the forming cup^69^ have also been proposed to contribute to optic cup invagination. Most of the cell rearrangements accompanying both optic vesicle evagination and optic cup invagination display a lateral component and could contribute to the lateral movements of OP cells. However, our analysis shows that the lateral displacement of *rx3:gfp*^*+*^ neural retina progenitors is more important after Tref (the biological reference time we used to synchronize the embryos, which corresponds to an invagination angle of 120°, see methods) when invagination rearrangements dominate. In addition, the mediolateral correlation coefficient between the eye and the OP was systematically higher after Tref than before Tref (Figure S2D). Moreover, degrading ECM from 16-18s, i.e. from the beginning of the invagination and after evagination, was sufficient to perturb lateral movements in the OP. Altogether these results suggest that optic cup invagination is the main actuator of OP’s shape change along the mediolateral axis. We hypothesise that the bending of the optic cup during invagination generates laterally oriented traction forces on OP cells, resulting in their lateral movements and the elongation of their axons. Additional experiments are required to identify the precise mechanism (apical constriction and cell compaction in the centre of the retina, rim movements of retinal cells, or spreading of the RPE over the cup) which generates these traction forces.

How is force transmitted between the eye and the OP? We observed the presence of NCCs and ECM at the interface between the two tissues. Interactions between NCCs and cranial placodes are crucial for the development of both cell populations in other contexts^25,70,71^. Using the *foxd3* mutant, in which cranial NCC migration is affected but eye morphogenesis is normal^46^, we demonstrate that the presence of NCCs is not required at the eye/OP interface to mediate OP lateral movements. On the contrary, degrading the ECM, which results in a physical uncoupling of the eye and the OP, clearly perturbs lateral movements in the OP. This strongly suggests that the ECM propagates the traction forces exerted by the eye to the OP. We showed the localisation of Laminin and Fibronectin at the tissue interface, but other ECM components are known to be also present in this area at similar stages^46^. Future experiments will determine whether the intertissue ECM meshwork as a whole, or a specific ECM component, transmits the forces from the eye to the OP.

It is increasingly evident that the biomechanics of morphogenesis must be understood not only in the scope of isolated tissues, but also in the framework of interacting tissues^32,33^. For instance, a recent study explained why elongation rates of paraxial and axial tissues are similar during elongation of the body axis elongation in chicken embryos. The compression of axial tissues by the flanking mesoderm leads to their elongation wich in turn promotes the mesoderm elongation too. It is tempting to imagine that such mechanical feedback loop involving adjacent tissues is a general mechanism of morphogenesis orchestration^72^. In Drosophila, endoderm invagination induces a tensile stretch contributing to germband extension^60,73^, but the mechanism behind such stress transmission remains elusive. ECM-mediated differential friction between tissues is essential for myotomes to acquire a chevron shape in zebrafish^65^. In addition, Fibronectin-mediated intertissue adhesion transmits shear forces between the neural tube and the paraxial mesoderm, which ensures symmetric morphogenesis of the zebrafish spinal cord^74^. Direct cell/cell contacts mediated by cadherins have been shown to transmit forces between adjacent tissues during gastrulation morphogenetic movements^75,76^ and the elongation of *C. elegans* embryos^77^. Our study adds to this emerging field by bringing new insight on how the building of neuronal circuits requires intertissue mechanical interaction, and presents an original example of a tissue elongation process resulting from the traction by an adjacent tissue which is connected by ECM.

Our results suggest a scenario in which the eye pulls on OP neuronal cells bodies, which stretches the anchored axonal protrusions that, in turn, grow in response to that tension. The retrograde axon extension in the OP could thus be seen as an *in vivo* example of stretch-induced or towed growth, in which axons grow in response to extrinsic tension without motile growth cones. This process has been hypothesised long ago by Paul Weiss to occur widely during development and body growth^78^, but has been mostly studied *in vitro* so far (for reviews see references^4,79,80^). OP morphogenesis thus represents a relevant *in vivo* model to investigate the mechanotransduction mechanisms by which neurons sense and transduce extrinsic forces into axon elongation, and in particular where and how novel material (membrane, cytoskeleton) is added along the axon shaft to accommodate stretch-induced growth. Mechanotransduction pathways can also drive changes in cell fate^81–83^ and could, in the OP, influence neurogenesis and neuronal differentiation, which are concomitant with OP morphogenesis^27,29,37,84^.

## Methods

### Fish strains

Wild-type, transgenic and mutant zebrafish embryos were obtained by natural spawning. In the text, the developmental times in hpf indicate hours post-fertilization at 28°C. To obtain the 12s stage, embryos were collected at 10 am, incubated for 2h at 28°C before being placed overnight in a 23°C incubator. 12s corresponds to 14 hpf and 24s to 21 hpf. The OP was visualised with the *Tg(8*.*4neurog1:gfp)* line^36^, referred to as *ngn1:gfp* in the text. The eye was visualised with the *Tg(rx3:GFP)* line^45^, referred to as *rx3:gfp* in the text. The *Tg(−8*.*0cldnb:LY-EGFP)* line, referred to as *cldnb:lyn-gfp*, was used to label the membranes of the neural retina and the retinal pigmented epithelium, as well as all cells of the OP^62,63^. The *rx3*^*s399*^ mutant^50^ (*chokh, chk*^*s399*^), referred to as *rx3* mutant, was used to examine OP morphogenesis in the absence of eye evagination. The *foxd3*^*zdf10*^ mutant^85^, referred to as *foxd3* mutant, was provided by the ZIRC Oregon and used to analyse OP morphogenesis when cranial NCC migration is perturbed. All our experiments were made in agreement with the European Directive 210/63/EU on the protection of animals used for scientific purposes, and the French application decree ‘Décret 2013-118’. The fish facility has been approved by the French ‘Service for animal protection and health’, with the approval number A-75-05-25.

### Genotyping

Homozygous *rx3*^-/-^ embryos were identified by their morphology/phenotype as the absence of eyes was easily detectable from 8-10s. For maintaining the line, the *rx3* locus was amplified from genomic DNA of adult fish using the 5’-TTATGCAGGAGTTTGTAGG-3’ (forward) and 5’-TAGTAGCCTATACTTCTCC-3’ (reverse) primers. The BspeI restriction enzyme (NEB, R0540S) cuts the wild type allele, giving rise to two fragments of 172bp and 159bp, but not the *rx3*^*s399*^ allele (331bp). The *foxd3*^*zdf10*^ allele was genotyped with the CAPS (Cleaved Amplified Polymorphic Sequences) technique^86^, using the SspI restriction enzyme (NEB, R132S), as described in^46^.

### mRNA injections

mRNAs were synthesised from linearised pCS2 vectors using the mMESSAGE mMACHINE SP6 transcription kit (Ambion). The following amounts of mRNA were injected into one-cell stage embryos: 80-100 pg for H2B-RFP, and 100 pg for mbCherry (membrane Cherry)^87^.

### Immunostainings

For immunostaining, embryos were fixed in 4% paraformaldehyde (PFA, in PBS), blocked in 5% goat serum, 1% bovine serum albumin and 0.3% triton in PBS for 2-3h at room temperature and incubated overnight at 4°C with primary and secondary antibodies. The following primary antibodies were used: anti-acetylated-tubulin (mouse, 1/500, 6-11B-1 clone, T6793, Sigma), anti-Laminin (rabbit, 1/100, L-9393, Sigma), anti-Fibronectin (rabbit, 1/100, F3648, Sigma).

### *In situ* hybridisation

Partial cDNA sequences for the NCC *crestin* marker were amplified by PCR using the 5’-AAGCCCTCGAAACTCACCTG-3’ (forward) and 5’-CCACTTGATTCCCACGAGCT-3’ (reverse) primers. PCR products were subcloned in pGEM-T-easy (Promega) and sequenced. The Digoxigenin(DIG)-labeled riboprobe was synthetized from PCR templates. Embryos were fixed in 4% PFA in PBS and stored in methanol at − 20 °C. Embryos stored in methanol were rehydrated in a methanol/PBS series, permeabilized 1min30s with proteinase K (10 mg/ml), pre-hybridized, and hybridized overnight at 65 °C in hybridization mixture (50% formamide, 5 X standard saline citrate (SSC), 0.1% Tween 20, 100 µg/lateral heparin, 100 µg/ml tRNA in water). The embryos were subjected to a series of washes in 50% SSC/formamide and SSC/PBST, and were then incubated in the blocking solution (0.2% Tween 20, 0.2% Triton X-100, 2% sheep serum in PBST) for one hour and overnight at 4 °C with alkaline phosphatase-conjugated anti-DIG antibodies (Roche) diluted at 1:4000 in the blocking solution. Embryos were then washed in PBST, soaked in staining buffer (TMN: 0,1M NaCl, 0,1M Tris-HCl, pH 9.5, 0.1% Tween 20 in water) and incubated in NBT/BCIP (nitroblue tetrazolium/5-bromo-4-chloro-3-indolyl phosphate) solution (Roche).

### Laser ablation of cells

Laser ablation of anterior and posterior converging cells was performed in OPs of 14-16s *ngn1:gfp* embryos injected with H2B-RFP mRNA to label the nuclei. Embryos were mounted in 0.5% low melting agarose in 1X E3 medium, and 10-20 cells in each anteroposterior extremity were ablated using a Leica TCS SP5 MPII upright multiphoton microscope with a ×25 objective (numerical aperture (NA): 0.95), laser at 790 nm). Two successive ablations were performed when the initial ablation did not generate any cell debris.

### Stress estimation at the OP periphery using laser ablation

Stress at the OP periphery was assessed by performing laser ablation of a line of cells. These supracellular laser ablations were performed in OPs of 14-18s *ngn1:gfp* embryos injected with mbCherry mRNA to label the membranes. Embryos were mounted in 0.5% low melting agarose in 1X E3 medium in Ibidi dishes (81158) and imaged using an inverted laser-scanning microscope (LSM 880 NLO, Carl Zeiss) equipped with a 63x oil objective (1.4 DICII PL APO, Zeiss). The cuts (about 20-25 µm long and 2-3 µm large) were performed in anterior, lateral and posterior borders of the *ngn1:gfp*^+^ cell cluster as depicted in Figure 5A. Ablations were performed using a Ti:Sapphire laser (Mai Tai, DeepSee, Spectra Physics) at 790 nm with <100 fs pulses, using a 80 MHz repetition rate, a 100% power and a number of iterations ranging between 5 and 10. Images were acquired at a frame rate of 0.46 s after ablation (the ablation itself lasted about 0.2 s). The initial velocity of wound margin retraction after ablation measures the stress-to-viscosity ratio within the tissue. To estimate the tension prior ablation, we thus measured the tissue flow by particle image velocimetry using the MatPIV toolbox for Matlab (Mathworks, US). The window size was set to 32 pixels (∼ 8 µm), with an overlap of 0.5. The 2D velocity field was measured between pre-cut and 5 frames after ablation (typically 2-3 s after ablation) in two rectangles of 10 µm wide adjacent and parallel to the line of ablation, as shown in Figure 5B. The “recoil velocity” was then obtained by summing the absolute values of the two average outward velocities (component orthogonal to the line of ablation) from each side of the ablation.

### Live imaging

Embryos were dechorionated manually and mounted at 11-12s in 0.5% low melting agarose in E3 medium, in order to obtain a frontal or a dorsal view of the head. Movies were recorded at the temperature of the imaging facility room (21-22°C) except for the collagenases-injection experiment (see below), on a Leica TCS SP5 AOBS upright confocal microscope or a Leica TCS SP5 MPII upright multiphoton microscope using ×25 (numerical aperture (NA) 0.95) or ×63 (NA 0.9) water lenses. At 21-22 °C, it takes around 800 min for zebrafish embryos to develop from 12s to 24 s stages. The Δt between each frame was 5 min for the analysis of cell movements in wild type embryos (Figures 2, S2) and 10 min for all the other live imaging experiments.

### Injection of collagenases

Embryos were injected at 1-cell stage with H2B:RFP mRNA, dechorionated manually at around 10s and mounted in 0.5% low melting agarose in E3 medium in order to obtain a dorsal view of the head. A collagenase mix was prepared with collagenase II (17101-015, Gibco) at 1000U/mL, collagenase IV (17104-019, Gibco) at 1000U/mL and Rhodamine Dextran (D1817, Invitrogen) at 5mg/mL diluted in E3 medium. Embryos were injected at 16-18s in the head region, close to the eye/OP interface, with a manual microinjector (CellTram Oil, Eppendorf). After injection, embryos were imaged using a Leica TCS SP5 MPII upright multiphoton microscope and ×25 (numerical aperture (NA) 0.95) water lens. Movies started around 30 minutes after the injection and lasted between 4 and 5 hours at 28°C. Immunostainings for Laminin were conducted on other injected embryos, kept at 28°C after injection and fixed at 24s.

### Image analysis

#### Quantification of OP volume, cell number and dimensions

To measure the volume of the OP *ngn1:gfp*^*+*^ cluster, we manually cropped the OP using ImageJ, segmented the *ngn1:gfp*^*+*^ cluster using median filtering and thresholding, and measured the volume of the object using the 3D ImageJ suite^88^. 3D counting of nuclei was achieved on the DAPI channel with the 3D ImageJ suite^88^ and the 3D Object Counter plugin^89^. To quantify the dimension of the OP, the *ngn1:gfp*^*+*^ segmented object was fitted with a 3D ellipsoid (see Figure 3B). A rotation of the image was applied so that the length of the ellipsoid 3D bounding box aligns with the orientation of the brain/placode interface. The lateral and anteroposterior dimensions of the OP correspond respectively to the X and Y dimensions of the 2D bounding box of the central z-slice of the ellipsoid, and the dorsoventral dimension corresponds to the Z dimension of the 3D bounding box of the whole ellipsoid.

#### Quantification of axon length and twisting index

Individual acetyl-tubulin positive axons were segmented manually in 3D, on images of fixed embryos at 24s, using the multiple point tool of Image J. The X,Y,Z coordinates of the resulting points were recovered and used to calculate the total 3D length of the axons and their twisting index, defined as 1 -(total 3D axon length/distance between the most proximal and the most distal point of the axon).

#### Manual cell tracking

Individual cells from the OP, the eye, the brain or the skin (periderm) were tracked in 3D using the Manual Tracking plugin in ImageJ. Tracked cells from the brain and the OP expressed *ngn1:gfp* at least at the end of the movie. Tracked cells from the eye expressed *rx3:gfp* at least at the end of the movie. Mediolateral cell positions were rescaled depending on the brain midline position to remove the contribution of a potential general drift of the embryo. Plots representing cell tracks merged at their origin were produced with Microsoft Excel and 2D color coded trajectories were generated in Matlab (Mathworks, US).

#### Time registration or synchronization

In order to compare similar stages of optic cup morphogenesis in the 4 embryos we imaged for the correlation analysis (Figure 2), we measured the invagination angle of the optic cup (*θ*, Figure S2A) as a function of time. We observed that *θ* decreases as morphogenesis goes and systematically passes through an inflection point when *θ* is approximately equal to 120° (Figure S2B). We used this inflexion point as a common time-coordinate in order to time-synchronize the 4 embryos. We thus fitted the invagination angle using the following equation: 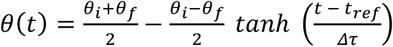 where *θ*_*i*_ and *θ*_*f*_ are the invagination angles at 12s and 30s respectively, *T*_ref_ the time corresponding to an invagination angle of 120° and **Δτ** the width of the *tanh*. We imposed the invagination angles *θ*_*I*_ = 180° and *θ*_*f*_ = 60°, in agreement with our observations. We thus shifted the curves of each movie in time so that their inflection points overlap for *θ*=120°. This enables us to set a common time window of 14h for the 4 embryos we analysed, depicted in light blue in Figure S2B.

#### Computation of correlation coefficients between pairs of cell tracks

The correlation coefficient *R* between two trajectories (*T*^(1)^, *T*^(2)^) is the covariance of the two trajectories normalized by the variance of each trajectory: 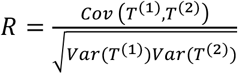. The 1D correlation coefficient for the mediolateral component (*x* component) is thus:

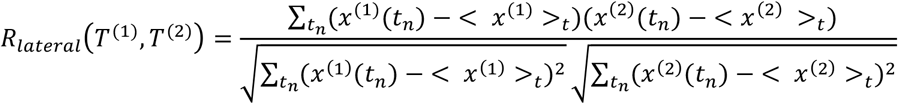

where <>_t_ is the temporal averaging. *R* lies in the range [-1; 1] and *R_2_* approximately indicates the percentage of the variation in one variable that can be explained by the variation in the other variable. Since the value of *R* is a function of the sample size, we consider the largest time-window common to all the embryos, which was 14h25min (depicted in light blue in Figure S2B), corresponding to 173 points.

### Statistical analysis

Graphs show mean ± s.e.m. (standard error of the mean), overlayed with all individual data points. The plots were generated with the GraphPad Prism software. For all figures, we checked for normality using Shapiro-Wilk normality tests, before performing parametric or non-parametric tests. The nature of the statistical test performed for each graph is indicated in the figure legends. The p values are corresponding to *p < 0.05, **p < 0.01, ***p < 0.001. No statistical method was used to estimate sample size and no randomisation was performed. Blinding was performed for the analysis of the supracellular laser ablation experiments.

**Figure S1. Additional results for the laser ablation of anteroposterior converging cells. A**. Images (dorsal view) of embryos 2 to 5 following the ablation (0 min) performed at 14-16s and 500 min later (24s). The OPs are surrounded by white lines. Magenta arrowheads show, when they are visible, the cellular debris resulting from cell ablation, in the anterior and/or posterior regions of the *ngn1:gfp*^+^ placodal clusters. Scale bars: 40 µm. **B**. Graphs showing the volumes of the ablated and control *ngn1:gfp*^+^ placodes for 3 time-points in embryos 2 to 5, t = 0 refers to the ablation. **C**. 2D-tracks of 5 to 7 OP central cells in the ablated and control placodes in embryos 2 to 5. The time is color-coded: light magenta at the starting of the trajectory (just after the ablation) towards dark magenta for the end of the track (500 min later). Scale bars: 10 µm.

**Figure S2. Quantitative analysis of cell movements: time-synchronisation of the embryos and additional results for the correlation analysis**. A. Confocal section of a *ngn1:gfp*; *rx3:gfp* double transgenic embryo injected with H2B:RFP mRNA, showing how the invagination angle of the optic cup (*θ*) was measured (shown here for embryo 1 at 24s, frontal view). B. *θ* decreases over time and systematically passes through an inflection point when *θ* is approximately equal to 120°. This inflexion point is used as a common time-coordinate (Tref) to time-synchronize the 4 embryos (see methods for more details). We shift the curves of each movie in time so that their reference times Tref overlap at *θ*=120°. This enables us to crop our movies in time to get a common time-window of 14h (represented in light blue on the graphs) to analyse cell movements for the 4 embryos. C. Intratissue mediolateral (ML) correlation coefficients. Each dot represents the average intratissue correlation, calculated from the tracking data of 5 to 7 cells per tissue (N = 4 embryos from 2 independent experiments). D. OP/eye mediolateral (ML) correlation coefficients for pairs of cells belonging to the placode and to the eye before and after Tref. E. Graphs showing the mediolateral (ML) displacement of OP, eye, brain and skin cells as a function of time (mean +/- sem from 5 to 7 cells per tissue). F. Correlation matrices presenting the mediolateral (ML) correlation coefficients between pairs of cell tracks. Each matrix represents the correlation coefficients for all pairs of cell tracks for a given pair of tissues in a given embryo, calculated from the tracking data of 5 to 7 cells per tissue (N = 4 embryos from 2 independent experiments). The average ML correlation coefficients (represented in Figure 2C-E for the embryo 1) for each tissue couple are indicated under the matrices.

**Figure S3. OP volume and number of cells are not affected in *rx3***^***-/-***^ **eyeless embryos**. Graphs showing the number of *ngn1:gfp*^+^ placodal cells (left) and the volume of the *ngn1:gfp*^+^ OP cluster (right) in *rx3*^*-/-*^ eyeless mutant and control embryos at 24s (N = 9 embryos from 3 independent experiments). Unpaired, two-tailed t test. Embryos from the same experiment are represented with similar markers (dots, triangles or squares).

**Figure S4. Additional results for the analysis of cell movements in *rx3***^***-/-***^ **eyeless embryos. A**. Graphs showing the total anteroposterior (AP) displacement of anterior, central and posterior OP cells, brain cells and skin cells, starting at 12s and during 700 min of time lapse in *rx3*^*-/-*^ eyeless mutants and control embryos (N = 3 control placodes and N = 5 mutant placodes from 3 independent experiments, mean calculated from 5 to 7 cells per tissue, unpaired, two-tailed t test). **B**. Graphs showing the total dorsoventral (DV) displacement of OP, brain and skin cells starting at 12s and during 700 min of time lapse in *rx3*^*-/-*^ eyeless mutants and control embryos (N = 3 control placodes and N = 5 mutant placodes from 3 independent experiments, mean calculated from 5 to 7 cells per tissue, unpaired, two-tailed t test).

**Figure S5. NCCs populate the eye/OP interface but are not essential for OP morphogenesis. A**. Image (dorsal view, 1 z-section) of a representative *cldnb:lyn-gfp; ngn1:gfp* double transgenic embryo fixed at 24s. The co-staining with DAPI (magenta) is shown on the left to visualise all nuclei. Note the absence of common cell/cell junctions between the eye and the OP. The white asterisks indicate the nuclei of gfp negative interstitial cells present in the gap between the eye and the OP. Scale bar: 20 µm. **B**. *In situ* hybridisation for the NCC marker *crestin*, in representative *foxd3*^*-/-*^ mutant (right) and *foxd3*^*+/+*^ control (left) embryos. Whole mount images and high magnifications of dorsal views on the head regions. **C**. Images (dorsal views) of representative placodes from a *ngn1:gfp; foxd3*^*-/-*^ mutant (right) and a *ngn1:gfp; foxd3*^*+/+*^ control sibling (left) at the end of OP coalescence (24s). The Laminin immunostaining (magenta) delineates brain, OP and eye tissues. White arrowheads indicate supposed interstitial NCC nuclei (DAPI labelling, cyan) between the basement membranes surrounding the tissues. **D**. Graphs showing, in *foxd3*^*+/+*^ controls and *foxd3*^*-/-*^ mutants, the area of the optic cup and its invagination angle calculated on a z-section passing through the centre of the lens (dorsal view), the OP volume, and mediolateral (ML), anteroposterior (AP), and dorsoventral (DV) dimensions of the OP at 24s, calculated as for *rx3*^*-/-*^ mutants in Figure 2 (N = 9 embryos from 3 independent experiments). Embryos from the same experiment are represented with similar markers (dots, triangles or squares). Unpaired, two-tailed t test except for the lateral dimension (no normal distribution) where a two-tailed Mann-Whitney test was performed.

**Figure S6. Additional results for the collagenases injection experiment. A**. Confocal image (dorsal view, 1 z-section) taken 20 min after injection of fluorescently labelled dextran (rhodamine, magenta) close to the eye/OP interface in a *ngn1:gfp; rx3:gfp* (green) embryo. The injected mix diffuses rapidly and fulfills all the space between tissues in the head region. **B**. Confocal images (dorsal views, 1 z-section) of embryos injected at 16s with collagenases and red dextran (right) or with dextran only (left), fixed at 24s and immunostained for Laminin. **C**. Graphs showing the mediolateral (ML) displacement of OP (magenta), eye (blue) and brain (green) cells as a function of time (mean +/- sem from 5 to 7 cells per tissue) for control and collagenases-injected embryos (experiments 2 and 3). To compensate for the drift often observed in injected embryos, the average displacement of skin cells was used for registration and removed from the tracking data. **D**. Correlation matrices displaying the intra-eye and the eye/OP mediolateral correlation coefficients, calculated from the tracking data of 5 to 7 cells per tissue (experiments 2 and 3). The average correlation coefficients for each tissue couple are indicated below the matrices.

## Video legends

**Video 1. OP morphogenesis upon laser ablation of anterior and posterior OP extremities**. Live imaging (dorsal view) of OP morphogenesis on a *ngn1:gfp* embryo in which laser ablation of anterior and posterior OP extremities was performed on the left placode at 16s (embryo 1). The contralateral, right placode is used as a control. In the first image, the ablated and control OPs are surrounded by white lines, and magenta arrowheads show cellular debris where cells were killed, in the anterior and posterior regions of the *ngn1:gfp*^+^ OP cluster. Maximum projection of a 84 µm stack.

**Video 2. Concomitant OP and eye morphogenesis**. Live imaging (frontal view) of OP morphogenesis from 12s to 24s on a *ngn1:gfp*; *rx3:gfp* double transgenic embryos embryo (embryo 1). The OP is outlined in magenta on the first and the last time points. Maximum projection of a 108 µm stack.

**Video 3. OP morphogenesis in *rx3***^***-/-***^ **eyeless and control embryos**. Live imaging (dorsal view) of OP morphogenesis from 12s to 24s on an eyeless *rx3*^*-/-*^ *ngn1:gfp* mutant (right) and a control *ngn1:gfp* embryo (left). The OPs are outlined in magenta on the last time point. Maximum projections of a 94 µm stack (control) and of a 56 µm stack (eyeless *rx3*^*-/-*^ mutant).

**Video 4. Example of a supracellular laser cut**. Recording of the tissue response after a line laser cut performed on the lateral border of the *ngn1:gfp*^*+*^ OP cluster in a control embryo expressing mbCherry to label the membranes (magenta). On the first time point, the region of ablation is shown in yellow (dorsal view, 1 z-section).

**Video 5. Confocal z-stacks on a live *cldnb:lyn-gfp* embryo**. Confocal z-stacks (dorsal view, z-step of 1 µm) on a live *cldnb:lyn-gfp* embryo at 14s (left) and 20s (right). In this transgenic line, a membrane-targeted version of gfp is expressed in the skin (periderm), the brain, in eye cells (both in the neural retina and the RPE) and in all cells of the OP. At both stages we observe the presence of a gfp negative gap between the eye and the OP (indicated by yellow arrowheads), showing the absence of shared cell/cell junctions between the two tissues.

**Video 6. 3D reconstruction on a live *cldnb:lyn-gfp* embryo**. 3D reconstruction of the z-stacks shown in Video 5. At both stages, 14s and 20s, we observe the presence of a clear gfp negative gap between the eye and the OP (indicated by yellow arrowheads), demonstrating the absence of shared intercellular junctions between the two tissues. Scale bar: 20 µm.

## Acknowledgements

This work was funded by the Agence Nationale pour la Recherche (ANR-17-CE13-0009-01 NEUROMECHANICS), the Centre National pour la Recherche Scientifique (CNRS, including the “Défi mécanobiologie” grant) and Sorbonne Université (including the Emergence 2016 grant, SU-16-R-EMR-11) and received support from the grants ANR-11-LABX-0038, ANR-10-IDEX-0001-02. M. Baraban was supported by a postdoctoral fellowship from the Fondation pour la Recherche Médicale (ARF201809006950). We also thank the imaging platform of the Institut de Biologie Paris-Seine (the facility is supported by CNRS, Sorbonne Université and the Conseil Régional Ile-de-France), the Cell and Tissue Imaging core facility (PICT IBiSA), Institut Curie, member of the French National Research Infrastructure France-BioImaging (ANR10-INBS-04), and the IBPS aquatic platform for fish care. We gratefully acknowledge Oxane Divaret for her investment during her two-month internship, and Pierre-Luc Bardet, Estelle Hirsinger, Julie Plastino, François Robin and Sylvie Schneider-Maunoury for insightful comments in reviewing the manuscript.

## Author contributions

Conceptualization: M.A.B. and I.B., Project administration: M.A.B. and I.B, Funding acquisition: M.A.B. and I.B, Methodology: M.A.B, I.B. and P.M., Software: I.B. and P.M., Investigation: P.M., G.G., M.B., K.P., M.C., Analysis: P.M., I.B, M.A.B., G.G., K.P., Supervision: M.A.B. and I.B, Writing: M.A.B, I.B. and P.M.

