## Supplementary figures and images for "Intertissue mechanical interactions shape the olfactory circuit in zebrafish"

### Supplemental Figures

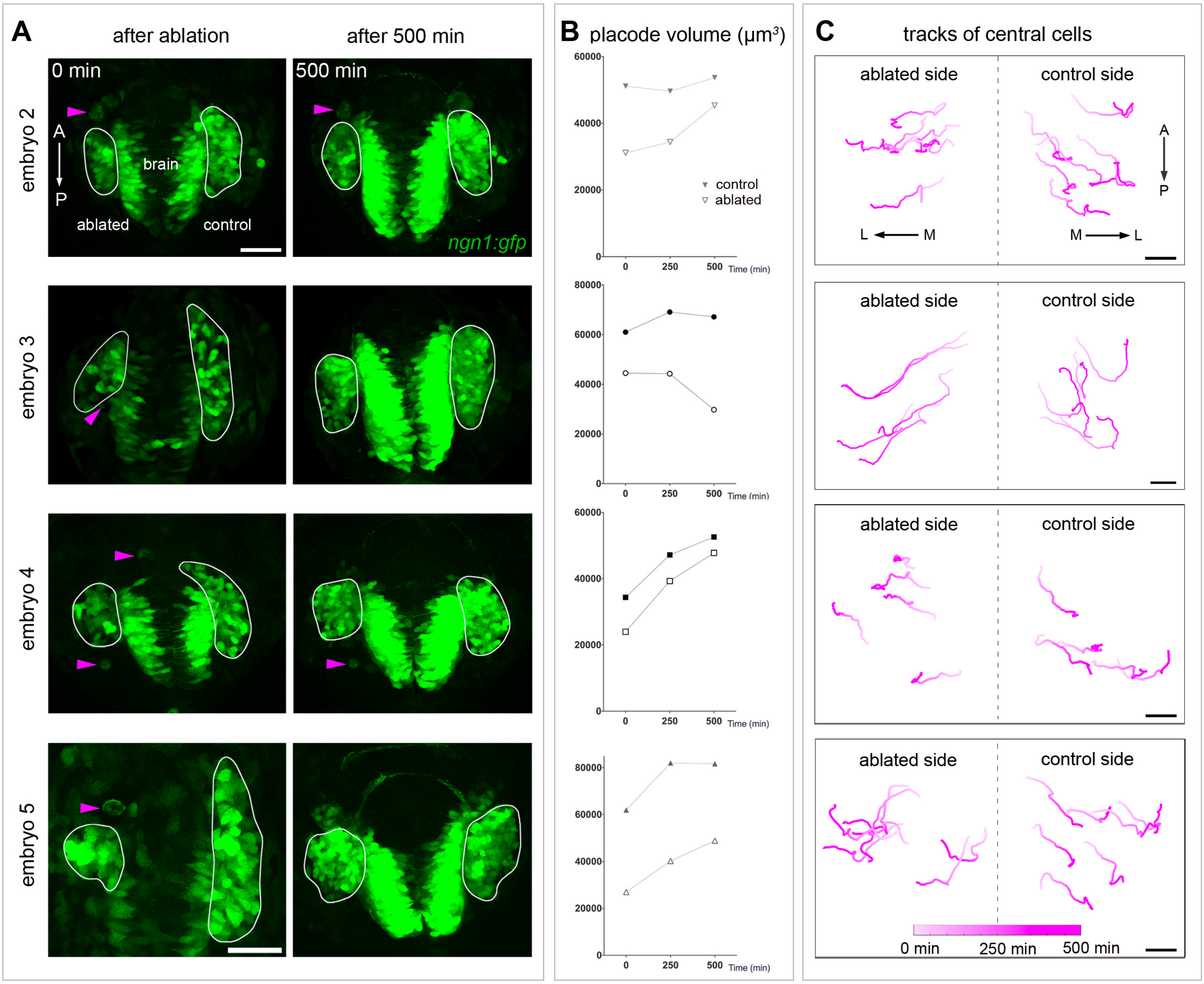

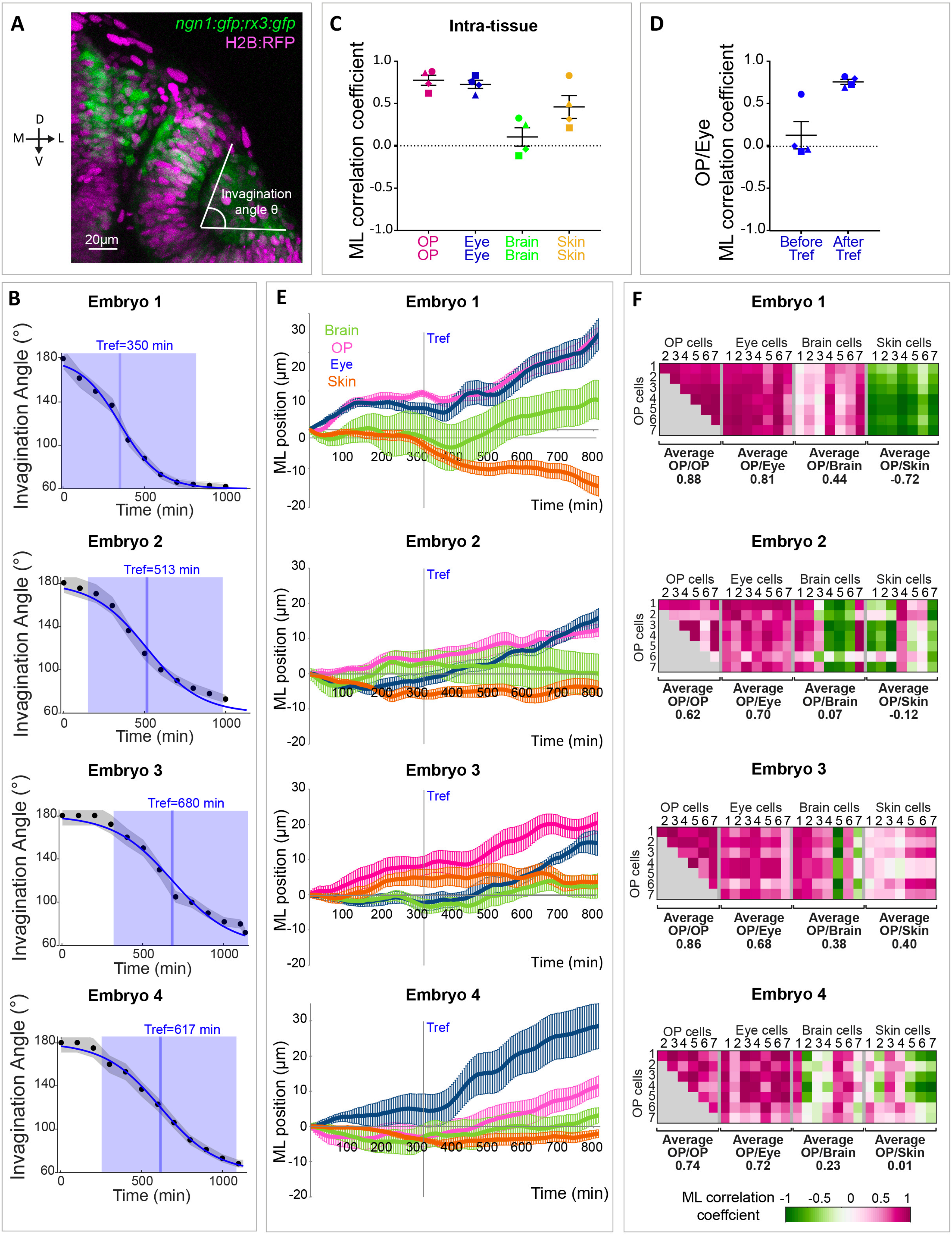

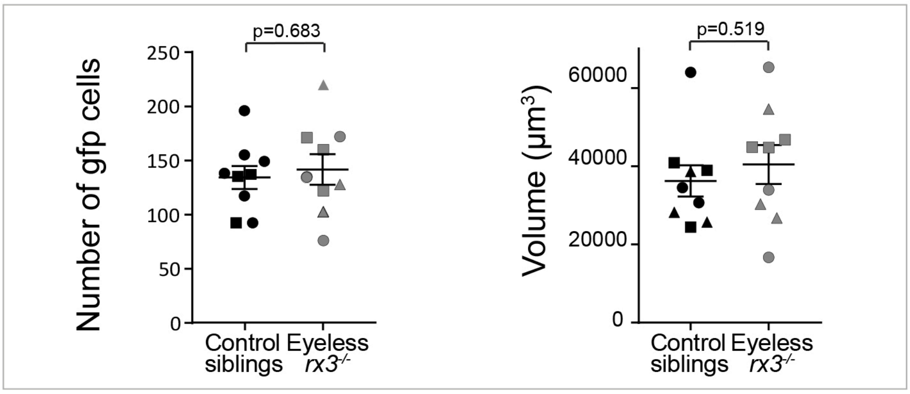

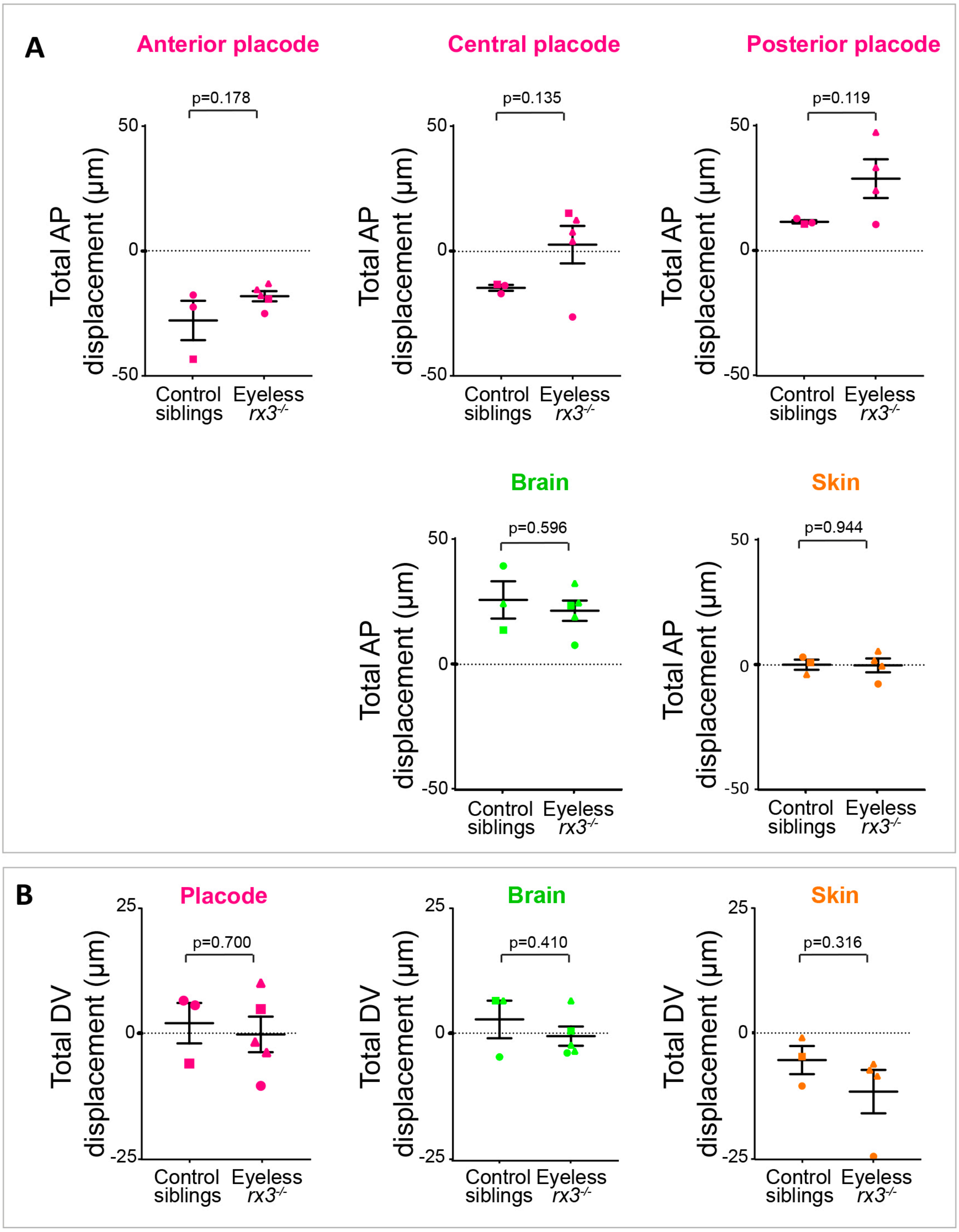

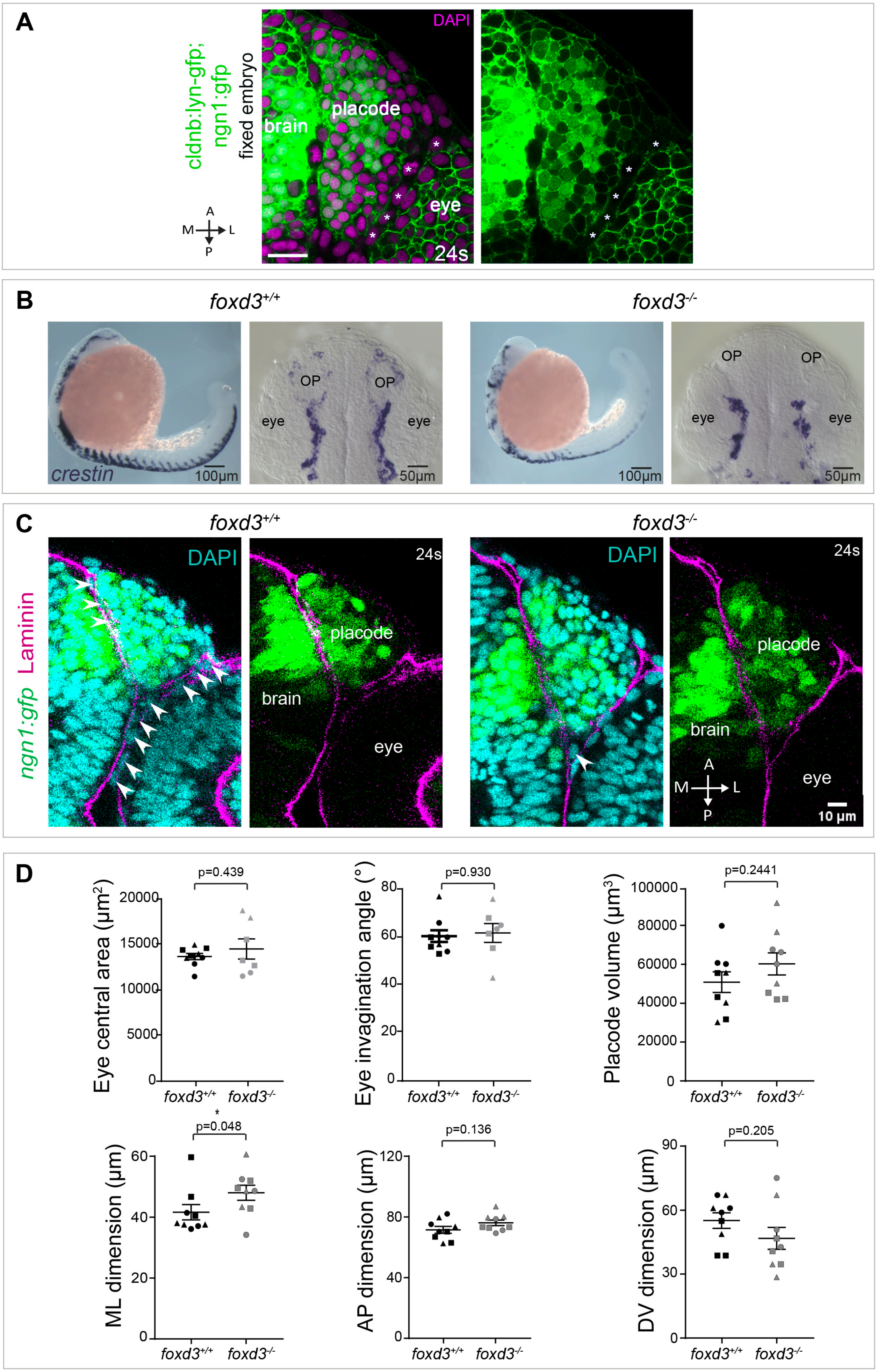

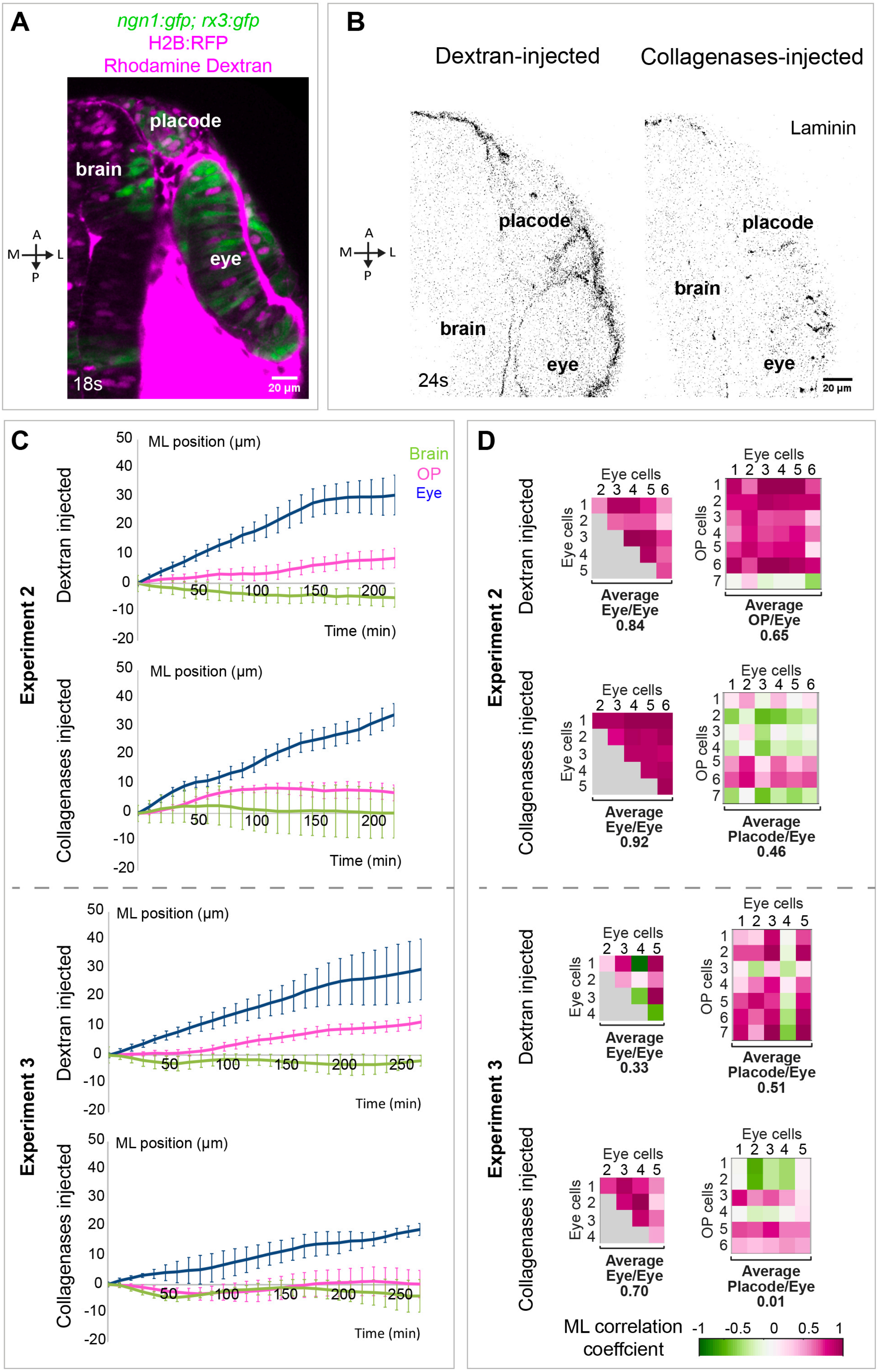
